# A genetic mouse model with postnatal *Nf1* and *p53* loss recapitulates the histology and transcriptome of human malignant peripheral nerve sheath tumor

**DOI:** 10.1101/2021.05.03.442481

**Authors:** Akira Inoue, Laura J. Janke, Brian L. Gudenas, Hongjian Jin, Yiping Fan, Joshua Paré, Michael R. Clay, Paul A. Northcott, Angela C. Hirbe, Xinwei Cao

## Abstract

**Background:** Malignant peripheral nerve sheath tumors (MPNST) are aggressive sarcomas. Somatic inactivation of *NF1* and cooperating tumor suppressors, including *CDKN2A/B*, PRC2, and p53, is found in most MPNST. Inactivation of the LATS1/2 kinases of the Hippo pathway was recently shown to cause tumors resembling MPNST histologically, although Hippo pathway mutations are rarely found in MPNST. Because existing genetically engineered mouse (GEM) models of MPNST do not recapitulate some of the key genetic features of human MPNST, we aimed to establish a mouse MPNST model that recapitulated the human disease genetically, histologically, and molecularly.

**Methods:** We combined two genetically modified alleles, an *Nf1*;*Trp53 cis*-conditional allele and an inducible *Plp-CreER* allele (NP-Plp), to model the somatic, possibly postnatal, mutational events in human MPNST. We also generated conditional *Lats1*;*Lats2* knockout mice. We performed histopathologic analysis of mouse MPNST models and transcriptomic comparison of mouse models and human nerve sheath tumors.

**Results:** Postnatal *Nf1;Trp53 cis*-deletion resulted in GEM-MPNST that was histologically more similar to human MPNST than the widely used germline *Nf1;Trp53 cis*-heterozygous (NPcis) model and showed partial loss of H3K27me3. At the transcriptome level, *Nf1;p53-*driven GEM-MPNST were distinct from *Lats*-driven GEM-MPNST and resembled human MPNST more closely than do *Lats-*driven tumors.

**Conclusions:** The NP-Plp model recapitulates human MPNST genetically, histologically, and molecularly.

**Key Points:**

1. Postnatal *Nf1;p53 cis*-deletion in NP-Plp mice results in tumors similar to MPNST.
2. The transcriptomes of *Nf1;p53-*driven and *Lats*-driven MPNST models are distinct.
3. NP-Plp model resembles human MPNST genetically, histologically, and molecularly.

**Importance of the Study:** Malignant peripheral nerve sheath tumors (MPNST) are aggressive sarcomas with a poor prognosis and limited treatment options. Existing genetically engineered mouse (GEM) models of MPNST do not recapitulate some of the key genetic features of human MPNST. To model the somatic, possibly postnatal, mutational events seen in MPNST patients, we generated a GEM-MPNST model by combining two genetically modified alleles, an *Nf1*;*Trp53 cis*-conditional allele and a *Plp-CreER* allele. Our histologic and transcriptomic analyses showed that this NP-Plp model resembles human MPNST genetically, histologically, and molecularly—more so than the widely used NPcis model and the recently published *Lats*-driven model. The NP-Plp model is genetically simple, making it easy to maintain and an ideal platform for preclinical studies. Given its tamoxifen-inducible nature, this model can be used to study the time/stage dependency of the tumorigenic potential of Schwann cells.

## Introduction

Malignant peripheral nerve sheath tumors (MPNST) are aggressive sarcomas arising from cells of the Schwann cell lineage. MPNST have a poor prognosis with limited treatment options. About half of MPNST occur in patients with the cancer predisposition syndrome neurofibromatosis type I (NF1), which is caused by germline loss-of-function (LOF) mutations in one copy of *NF1* (encoding neurofibromin). Other cases occur sporadically or after radiotherapy in the general population. Somatic mutations in *NF1* with co-occurring LOF alterations in *CDKN2A*, polycomb repressive complex 2 (PRC2) genes (*SUZ12* and *EED*), and/or *TP53* (*Trp53* in mice, encoding p53) are found in the vast majority of MPNST regardless of their etiology.^1^

Genetically engineered mouse (GEM) models of human diseases are useful for studying disease mechanisms and testing diagnostic/therapeutic approaches. Although several GEM models of MPNST have been published,^2,3^ they do not recapitulate some of the key genetic features of human MPNST. For example, the first and one of the most frequently used GEM-MPNST models harbors germline *cis*-heterozygous LOF mutations in *Nf1* and *Trp53* (NPcis mice),^4,5^ whereas MPNST patients do not carry germline *TP53* mutations, except for rare occurrences in Li-Fraumeni syndrome.^6^ Most, if not all, conditional GEM models use *Cre* lines that initiate gene deletion in embryonic Schwann cell lineage cells, such as *Dhh-Cre*,^7^ *Periostin-Cre*,^8^ and *GFAP-Cre*,^3,9^ although it is unclear whether somatic alterations in patients occur in Schwann cell precursors or immature Schwann cells during fetal development or in mature Schwann cells later in life. Injecting adenovirus expressing Cre or Cas9-gRNA into the sciatic nerve has the advantage of temporally controlled gene deletion;^3,10^ however, surgery is required to expose the nerve and may cause nerve injury.^11^ Although these existing GEM-MPNST models are useful for many studies, they are not ideal for addressing questions such as those regarding tumor cell-of-origin.

Most GEM-MPNST models were created based on the knowledge that nearly all MPNST harbor *NF1* mutations. Wu et al. reported that deleting *Lats1* and *Lats2* (*Lats1;2*) of the Hippo pathway in the Schwann cell lineage generated tumors histologically similar to MPNST.^12^ It was not shown, however, whether these *Lats*-driven GEM-MPNST resembled human MPNST or *Nf1*-driven GEM-MPNST at the transcriptome level.

Here, we used conditional alleles of *Nf1*^13^ and *Trp53*^14^ to generate a *cis*-conditional allele, [*Nf1;Trp53*]^*fl*^. Mice carrying [*Nf1;Trp53*]^*fl*^ and the inducible *Plp-CreER*,^15^ which targets multiple glial cell types in the peripheral nervous system,^16^ developed GEM-MPNST when [*Nf1;Trp53*]^*fl*^ was deleted postnatally. This new model recapitulated the histopathologic features of human MPNST better than the NPcis model. Concurrent with the study by Wu et al.,^12^ we independently generated *Lats*-driven GEM-MPNST models and found that the transcriptomes of *Nf1*-driven and *Lats*-driven GEM-MPNST were distinct. Further cross-species transcriptomic comparison suggests that *Nf1;p53*-driven GEM-MPNST models resemble human MPNST more closely than do *Lats*-driven GEM-MPNST models. Therefore, the NP-Plp model most closely recapitulates human MPNST genetically, histologically, and molecularly.

## Materials and Methods

### Animals

NPcis (Stock #008191), *Nf1*^*fl*^ (#017640), *Trp53*^*fl*^ (#008462), *Plp-CreER*^*T*^ (#005975), and *Nestin-Cre* (#003771) were obtained from The Jackson Laboratory. *Lats1*^*fl*^ and *Lats2*^*fl*^ were obtained from Randy L. Johnson.^17^ NPcis, *Nf1*^*fl*^, *Trp53*^*fl*^, and [*Nf1;Trp53*]^*fl*^ were maintained on the B6;129 mixed background obtained from Jackson, *Plp-CreER*^*T*^ and *Nestin-Cre* on the C57BL/6J background, and Lats-Nes and Lats-Plp on a mixed background.

Tamoxifen (Sigma-Aldrich T5648) was dissolved in corn oil. For P7 mice,15μl/g body weight of a 5mg/ml solution was injected intraperitoneally once daily for 2 consecutive days. For 4–8-week-old mice, 5μl/g body weight of a 20mg/ml solution was administered by oral gavage once daily for 3 consecutive days. All procedures were approved by the St. Jude Children’s Research Hospital Animal Care and Use Committee. Male and female mice were used in a ~1:1 ratio.

### Histology

Tissues were fixed in 10% formalin or 4% paraformaldehyde overnight. Tumors with adjacent bones were decalcified in 10% formic acid. Tissues were embedded in paraffin and sectioned at 4-μm or 6-μm thickness. Hematoxylin-eosin (H&E) staining was performed per routine methods. For immunostaining, slides were subjected to antigen retrieval at 95°C for 30 min in 10mM sodium citrate (pH 6.0) and incubated with primary antibody (Supplementary Table 1) overnight at 4°C. Images were acquired using a Keyence BZ-X700 or an Olympus BX46 microscope with an Olympus SC180 camera and processed in Photoshop. For Ki67 and H3K27me3 quantifications, each data point represents an individual tumor and is the average of measurements from three 182μm × 182μm regions per tumor.

### Western blot

Tumor and cell lysates were prepared using a bead homogenizer (Bertin) in 20 mM HEPES (pH 7.4), 150 mM NaCl, 2% SDS, and 5% glycerol supplemented with AEBSF and Halt protease and phosphatase inhibitors. Protein concentration was measured using BCA. Lysates were subjected to SDS-PAGE, probed with primary antibodies (Supplementary Table 1) and HRP-conjugated secondary antibodies.

### RNA isolation and sequencing

Mouse tumors were homogenized in TRIZOL using a bead homogenizer. Total RNA was isolated using the Direct-zol RNA Miniprep Kit (Zymo). Libraries were prepared using the TruSeq Stranded Total RNA Library Prep Kit (Illumina), analyzed for insert size distribution using the 4200 TapeStation D1000 ScreenTape Assay (Agilent), and sequenced on Illumina NovaSeq 6000, yielding 100 million 100-bp paired-end reads. Sequences are deposited in GEO (GSE172221).

### RNA-seq data analysis

Mouse and human sequences were mapped to the mm10 and hg38 genomes, respectively, with STAR aligner. Gene-level quantification was determined using RSEM and based on GENCODE M22 gene annotation. Non-coding and GENCODE level 3 genes were excluded. Differential expression was modeled using the voom method, available in the limma R package. Voom-normalized counts were analyzed using the lmFit and eBayes functions in limma. The false discovery rate (FDR) was estimated using the Benjamini-Hochberg method. Gene set enrichment analysis (GSEA) was performed using MSigDB gene sets as described.^18^ Single-sample GSEA (ssGSEA) was performed as described.^19^

### Cross-species transcriptome comparison

RNA-seq data from the *Lats;Dhh-Cr*e model (GSE99040), *Nf1* models (GSE148249), MPNST samples from The Cancer Genome Atlas (TCGA), and the MPNST and PNF cohort (GSE145064) were summarized to fragments per kilobase per million mapped fragments (FPKM) based on Ensembl genes (v93). These datasets were integrated with our mouse tumor samples and our (SJ collection) MPNST, PNF, and NF samples by filtering genes to mouse-human orthologues that were present in all datasets. Batch-effect was removed using ComBat^20^ to correct for batch and species differences. ssGSEA scores^19^ were derived using a list of orthologous gene sets created by retaining all gene sets from MSigDB that contain ≥80% mouse-human orthologues, ≥15 genes, and ≤400 genes.

To assess the resemblance of murine tumor models to human tumors in an unsupervised manner, we used uniform manifold approximation and projection (UMAP) based on the Pearson’s distance of the top 500 most varied gene sets across all murine models. A supervised analysis was performed by training a random forest classifier to predict whether human tumors were MPNST or PNF/NF based on the ssGSEA scores. The human tumor samples in the SJ collection and in GSE145064, comprising 34 MPNST and 33 PNF/NF samples, were split in a 70-30 ratio to generate the training and testing datasets. The top 20 most informative gene sets were used as features for training, which achieved a test set sensitivity of 0.825, specificity of 0.86, and accuracy of 0.80. This model was then used to predict the MPNST class probability for murine models and the independent test set of TCGA MPNST samples.

## Results

### Postnatal somatic *Nf1;Trp53* loss results in malignant peripheral nerve sheath tumors

Mouse *Nf1* and *Trp53* both locate on chromosome 11. To mimic somatic loss of *NF1* and *TP53* in MPNST patients, we generated a chromosome 11 harboring the conditional alleles of both *Nf1* and *Trp53*, designated [*Nf1;Trp53*]^*fl*^, by crossing *Nf1*^*fl/+*^;*Trp53*^*fl/+*^ *trans*-heterozygous mice with wild-type (WT) mice and screening for progeny carrying both conditional alleles as a result of meiotic recombination that placed them on the same chromosome. Mice homozygous for this chromosome were healthy and fertile, suggesting that the chromosome integrity was not affected by the recombination event. To enable temporal control of [*Nf1;Trp53*]^*fl*^ deletion, we chose the tamoxifen-inducible *Cre* line, *Plp-CreER*, which targets multiple glial cell types in the peripheral nervous system.^16^ We then combined the [*Nf1;Trp53*]^*fl*^ and *Plp-CreER* alleles with a germline WT or a null allele of *Nf1* (*Nf1−*), generating [*Nf1;Trp53*]^*fl/+*^;*Plp-CreER* and [*Nf1;Trp53*]^*fl/Nf1−*^;*Plp-CreER* mice, respectively, to mimic the genetics of sporadic and NF1-associated MPNST (Figure 1A).

**Figure 1.**
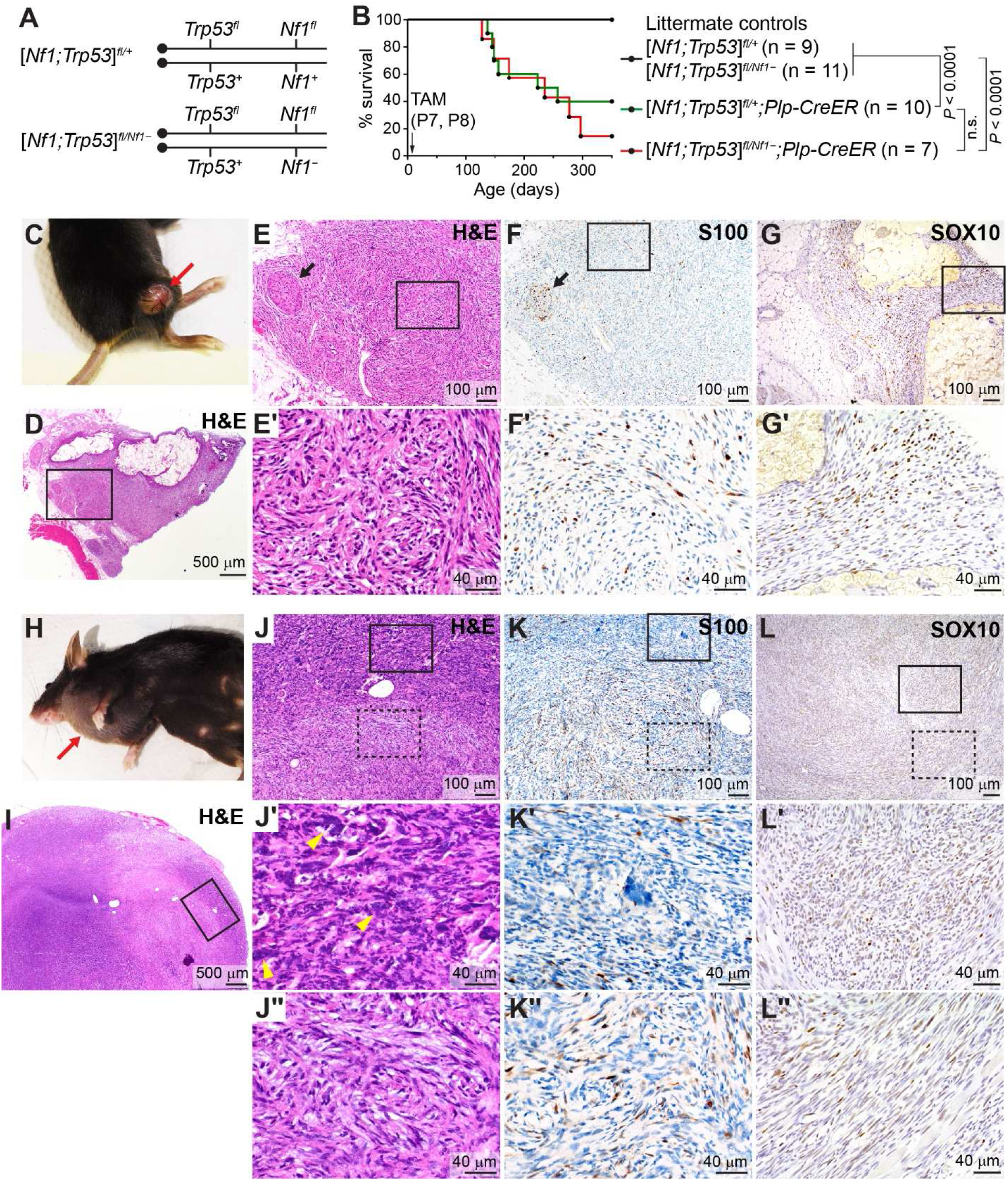
Postnatal somatic *Nf1;Trp53* loss results in malignant peripheral nerve sheath tumors. (A) Nomenclature of *Nf1;Trp53* genotypes and corresponding genetic alterations on mouse chromosome 11. (B) Kaplan-Meier survival curves of tamoxifen (TAM)-treated mice. Log-rank Mantel-Cox test. P, postnatal day. (C-G) Example of a dermal tumor, showing a [*Nf1;Trp53*]^*fl/+*^;*Plp-CreER* mouse with a ulcerated dermal tumor (C, red arrow) and histology of a grade II tumor stained with H&E (D, E) and S100 and SOX10 antibodies (F, G, brown signal). Boxed area in D is enlarged in E. Boxed areas in E, F, G are enlarged in E’, F’, G’. Arrows in E and F indicate a nerve-like structure. (H-L) Example of a subcutaneous tumor, showing an [*Nf1;Trp53*]^*fl/Nf1−*^;*Plp-CreER* mouse with a tumor in the axillary region of the left forearm (H, red arrow) and a grade III GEM-MPNST stained with H&E (I, J) and S100 and SOX10 antibodies (K, L). Boxed area in I is enlarged in J, showing regional heterogeneity. Solid-boxed area in J is enlarged in J’, showing a poorly differentiated area. Yellow arrowheads indicate examples of nuclear atypia. Dashed-boxed area in J is enlarged in J’’, showing spindle cells forming interlacing fascicles and storiform formations. Solid-boxed areas in K and L are enlarged in K’ and L’, and dashed-boxed areas are enlarged in K’’ and L’’.

As an initial test, we administered tamoxifen on postnatal day (P) 7 and P8, when immature Schwann cells proliferate, sort axons, and myelinate.^21^ Six of 10 [*Nf1;Trp53*]^*fl/+*^;*Plp-CreER* and six of seven [*Nf1;Trp53*]^*fl/Nf1−*^;*Plp-CreER* mice developed one or two palpable masses at 4–9 months of age (Figure 1B). Six of the 16 masses were firm and grew relatively slowly, and the mice had to be euthanized within 4 weeks of tumor appearance because of tumor ulceration (Figure 1C). These tumors were all in the skin. Of the other 10 masses, six were subcutaneous, associated with skeletal muscles, and four were in the abdominal cavity. These masses were soft, grew rapidly, and mice had to be euthanized within 1 week of tumor appearance because of tumor size (Figure 1H).

We performed histopathologic analysis of these tumors according to criteria established for mouse models of PNST^22^ and criteria used for human nerve sheath tumors.^23^ All five dermal tumors analyzed exhibited features typical of GEM-MPNST, comprising spindle cells forming interlacing fascicles and storiform formations with high cellularity and occasional pleomorphism and nuclear atypia (Figure 1D, E). Two were diagnosed as grade II GEM-MPNST based on their low mitotic rate and low-to-moderate levels of Ki67 staining. The remaining three were diagnosed as grade III GEM-MPNST based on the presence of regions with a higher mitotic rate. The eight fast-growing, internal tumors analyzed were all diagnosed as grade III GEM-MPNST. These tumors had growth patterns similar to those of the grade II GEM-MPNST but had more frequent pleomorphism and nuclear atypia, a high mitotic rate, and areas of necrosis. They more frequently contained poorly differentiated areas composed of smaller cells with a high nucleus-to-cytoplasm ratio (Figure 1I, J). Both grade II and III tumors showed multifocal nuclear and occasionally cytoplasmic S100 immunolabeling (Figure 1F, K). The diagnosis of GEM-MPNST was further supported by moderately positive immunolabeling for SOX10 in two of four tumors tested (Figure 1G, L) and widespread positive Nestin immunolabeling in all tumors (data not shown).

Thus, postnatal somatic *Nf1*;*Trp53* loss resulted in GEM-MPNST that resembled human MPNST. Because we detected no difference in tumor onset, histology, or survival between [*Nf1;Trp53*]^*fl/+*^;*Plp-CreER* and [*Nf1;Trp53*]^*fl/Nf1−*^;*Plp-CreER* mice—which might be caused by the small number of mice analyzed—we designated these mice/tumors collectively as NP-Plp mice/tumors.

### NP-Plp GEM-MPNST is histologically distinct from NPcis GEM-MPNST

Next, we compared our NP-Plp model to the NPcis model, one of the most frequently used MPNST models. Twelve of the 13 NPcis mice analyzed developed palpable subcutaneous tumors at 4–6 months of age (Figure 2A). These tumors were soft and fast-growing, and mice had to be euthanized within 1 week of tumor appearance because of tumor size. Gross dissection revealed that two mice harbored an additional soft tumor, one in the abdomen and one in the thoracic cavity. The remaining mouse developed three firm, slow-growing dermal tumors at 4 months of age and was euthanized 1 month after tumor appearance because of ulceration (Figure 2F).

**Figure 2.**
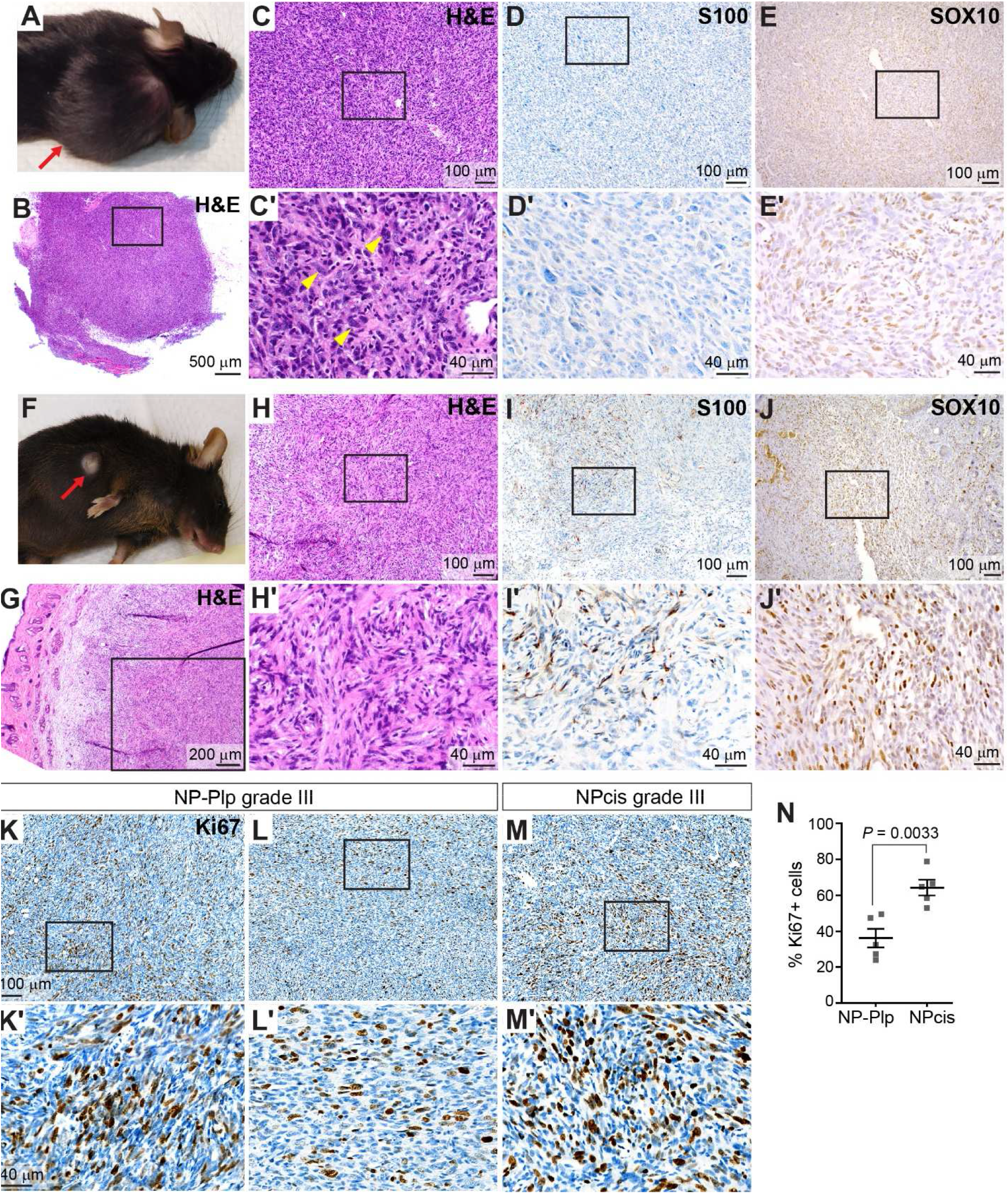
Histopathologic comparison of NP-Plp and NPcis GEM-MPNST. (A-E) Example of a subcutaneous NPcis grade III GEM-MPNST, showing an NPcis mouse with a tumor over the right shoulder (A, red arrow) and H&E staining (B, C) and S100 and SOX10 immunostaining (D, E). Boxed area in B is enlarged in C. Boxed areas in C, D, E are enlarged in C’, D’, and E’. Yellow arrowheads in C’ indicate nuclear atypia. (F-J) Example of a dermal NPcis grade II GEM-MPNST, showing an NPcis mouse with a tumor (F, red arrow) and H&E staining (G, H) and S100 and SOX10 immunostaining (I, J). Boxed area in G is enlarged in H. Boxed areas in H, I, J are enlarged in H’, I’, and J’. (K-N) Ki67 immunostaining and quantification (mean±SEM) in grade III tumors. Unpaired, two-tailed *t* test.

Five of the seven soft, fast-growing tumors analyzed exhibited nearly identical histopathologic features (Figure 2B, C). These subcutaneous tumors were homogeneous, comprising small cells with mild spindling. The cells were more pleomorphic and undifferentiated and had a higher nucleus-to-cytoplasm ratio than is typical in NP-Plp tumors. The nuclei were often hyperchromatic and enlarged with marked atypia. The tumors had a high mitotic rate. Although the spindle-cell morphology and fascicular and storiform patterns typically found in MPNST were present only occasionally in these tumors and SOX10 immunolabeling was weak, most tumors showed occasional S100 signal along with Nestin immunolabeling (Figure 2D, E and data not shown), and were, therefore, diagnosed as grade III GEM-MPNST. The remaining two soft, fast-growing tumors were diagnosed as lymphoma and were excluded from further analyses. The two dermal tumors analyzed exhibited typical MPNST features (Figure 2G-J), including spindle cells forming interlacing fascicles and storiform patterns, had a low mitotic rate, and were diagnosed as grade II GEM-MPNST. Quantification of Ki67 signals in grade III NP-Plp and NPcis GEM-MPNST showed the proliferation rate to be significantly higher in NPcis tumors than in NP-Plp tumors (Figure 2K-N).

### LATS1/2 loss in the Schwann cell lineage results in GEM-MPNST

While studying the role of LATS1/2 during nervous system development, we deleted *Lats1;2* with a *Nestin-Cre* line that targets progenitor cells in the central and peripheral nervous system, and serendipitously found that mice lacking three alleles of *Lats1;2*, *Lats*^*fl/fl*^;*Lats2*^*fl/+*^;*Nestin-Cre* and *Lats1*^*fl/+*^;*Lats2*^*fl/fl*^;*Nestin-Cre* (referred to as Lats-Nes mice), all developed tumors at 4–6 months of age. Tumors were most frequently found in the skin, with each mouse typically having multiple firm, slow-growing tumors, and the mice had to be euthanized 2–4 months after tumor appearance, usually because of tumor ulceration and occasionally because of limb paralysis (Figure 3A). Gross dissection revealed additional firm, internal tumors that were often associated with nerves and neural ganglia. Histopathologic analysis revealed typical features of GEM-MPNST. All paraspinal and nerve-associated tumors were diagnosed as grade III GEM-MPNST, whereas some dermal tumors were diagnosed as grade III and others as grade II GEM-MPNST (Figure 3B, C). To confirm that GEM-MPNST formation was due to *Lats1;2* loss in the peripheral glial lineage, we used *Plp-CreER* to delete *Lats1;2*. Tamoxifen administration to 1–2-month-old *Lats1*^*fl/+*^;*Lats2*^*fl/fl*^;*Plp-CreER* (Lats-Plp) mice resulted in dozens of firm, slow-growing dermal tumors in each mouse and occasional internal tumors with 100% penetrance (Figure 3D). All Lats-Plp tumors were grade II GEM-MPNST (Figure 3E). While our study was underway, Wu et al. reported similar findings when three alleles of *Lats1;2* were deleted with *Dhh-Cre* (Lats-Dhh) or *Plp-CreER*.^12^

**Figure 3.**
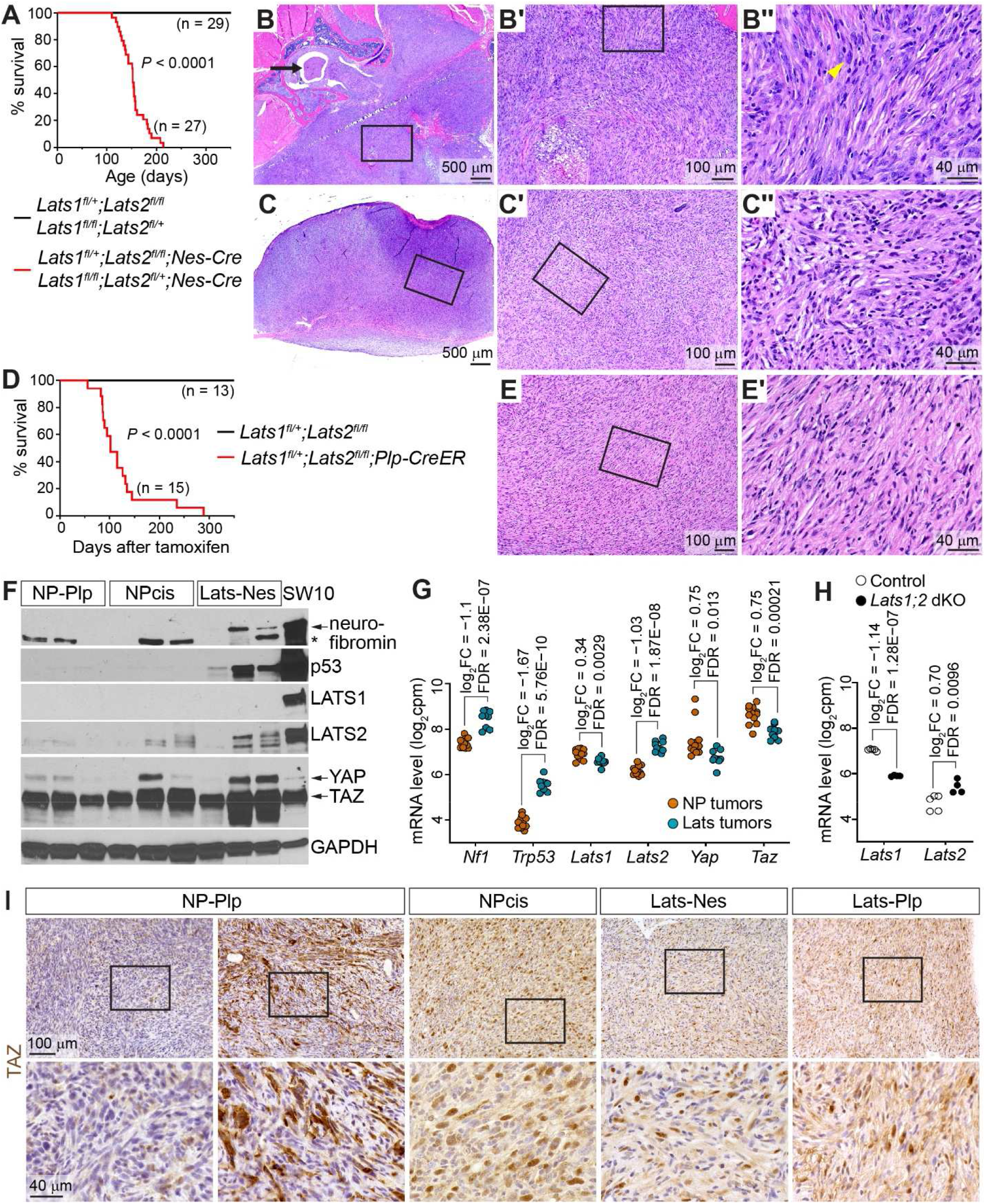
LATS1/2 loss causes GEM-MPNST formation and comparison to *Nf1;p53*-driven GEM-MPNST. (A) Survival curves of Lats-Nes mice and littermate controls. Log-rank Mantel-Cox test. (B) A large infiltrative grade III paraspinal tumor stained with H&E. Boxed areas in B and B’ are enlarged in B’ and B’’, respectively. Arrow in B indicates the spinal cord. Yellow arrowhead in B’’ indicates nuclear atypia. (C) A grade III dermal tumor stained with H&E. (D) Survival curves of Lats-Plp mice and littermate controls. Log-rank Mantel-Cox test. (E) A grade II Lats-Plp GEM-MPNST stained with H&E. (F) Western blot analysis of key oncogenic drivers. The immortalized Schwann cell line SW10 was used as the positive control. * indicates a possible truncated neurofibromin product. All samples were grade III tumors inferred based on tumor location. 20 μg of tumor lysate or 5 μg of SW10 cell lysate was loaded per lane. (G) mRNA counts obtained from RNA-seq analysis of tumor samples and fold changes (FC) in NP (NP-Plp and NPcis) versus Lats (Lats-Nes and Lats-Plp) tumors. Cpm, counts per million. (H) *Lats1* and *Lats2* mRNA counts obtained from RNA-seq analysis of E12.5 *Lats1*^*fl/fl*^;*Lats2*^*fl/fl*^;*Nestin-Cre* double knockout (dKO) and control (no *Cre*) telencephalons. (I) TAZ immunostaining. Boxed areas in the upper-row images are enlarged in the lower row.

### Differential expression of driver tumor suppressors in *Nf1;p53*-driven and *Lats*-driven GEM-MPNST models

We compared the expression of the key oncogenic drivers in *Nf1;p53*-driven (NP) and *Lats*-driven (Lats) GEM-MPNST models. As expected from the genotype, neurofibromin and p53 proteins were undetectable in NP tumors but were present in at least some Lats tumors (Figure 3F). Surprisingly, LATS1/2 protein levels were very low in NP tumors, similar to those in Lats tumors in which the *Lats1;2* genes were deleted. LATS1/2 are key kinases of the Hippo pathway. They phosphorylate transcriptional coactivators YAP and TAZ (YAP/TAZ) and prevent them from entering the nucleus and activating transcription. Therefore, LATS1/2 loss was expected to cause YAP/TAZ activation. Indeed, NP and Lats tumors all contained high levels of TAZ protein and abundant nuclear TAZ immunosignal (Figure 3F, I).

We also examined the mRNA levels of these oncogenic drivers using our RNA sequencing (RNA-seq) data (see below). Consistent with the genotype, *Nf1* and *Trp53* mRNA levels were markedly lower in NP tumors than in Lats tumors (Figure 3G). However, *Lats1* mRNA levels were only slightly lower in Lats tumors than in NP tumors. The reason for this is unclear, as our previous RNA-seq data^24^ showed that *Lats1* mRNA levels were markedly reduced in *Lats1*^*fl/fl*^;*Lats2*^*fl/fl*^;*Nestin-Cre* double knockout (dKO) brains compared to control brains (Figure 3H). *Lats2* mRNA levels were higher in Lats tumors than in NP tumors, just as they were higher in *Lats1;2* dKO brains than in control brains (Figure 3G, H). This was probably because of the feedback activation of the *Lats2* gene by activated YAP/TAZ.^25^ *Yap* and *Taz* mRNA levels were similar in NP and Lats tumors. In summary, whereas neurofibromin and p53 proteins were absent in NP tumors but present in Lats tumors, NP and Lats tumors both lacked LATS1/2 proteins and contained high levels of TAZ.

### Partial H3K27me3 loss in NP-Plp and Lats GEM-MPNST, but not in NPcis GEM-MPNST

PRC2 catalyzes the di- and tri-methylation of histone H3 at lysine 27. Somatic LOF mutations in PRC2 components are common in MPNST, leading to H3K27me3 loss.^26–28^ Immunostaining revealed that both grade II and III NP-Plp tumors showed varying degrees of H3K27me3 loss, which often exhibited intratumor heterogeneity (Figure 4A). In contrast, H3K27me3 signal was retained nearly uniformly in grade II and III NPcis tumors. Lats-Nes and Lats-Plp tumors also showed varying degrees of H3K27me3 loss (Figure 4A, B).

**Figure 4.**
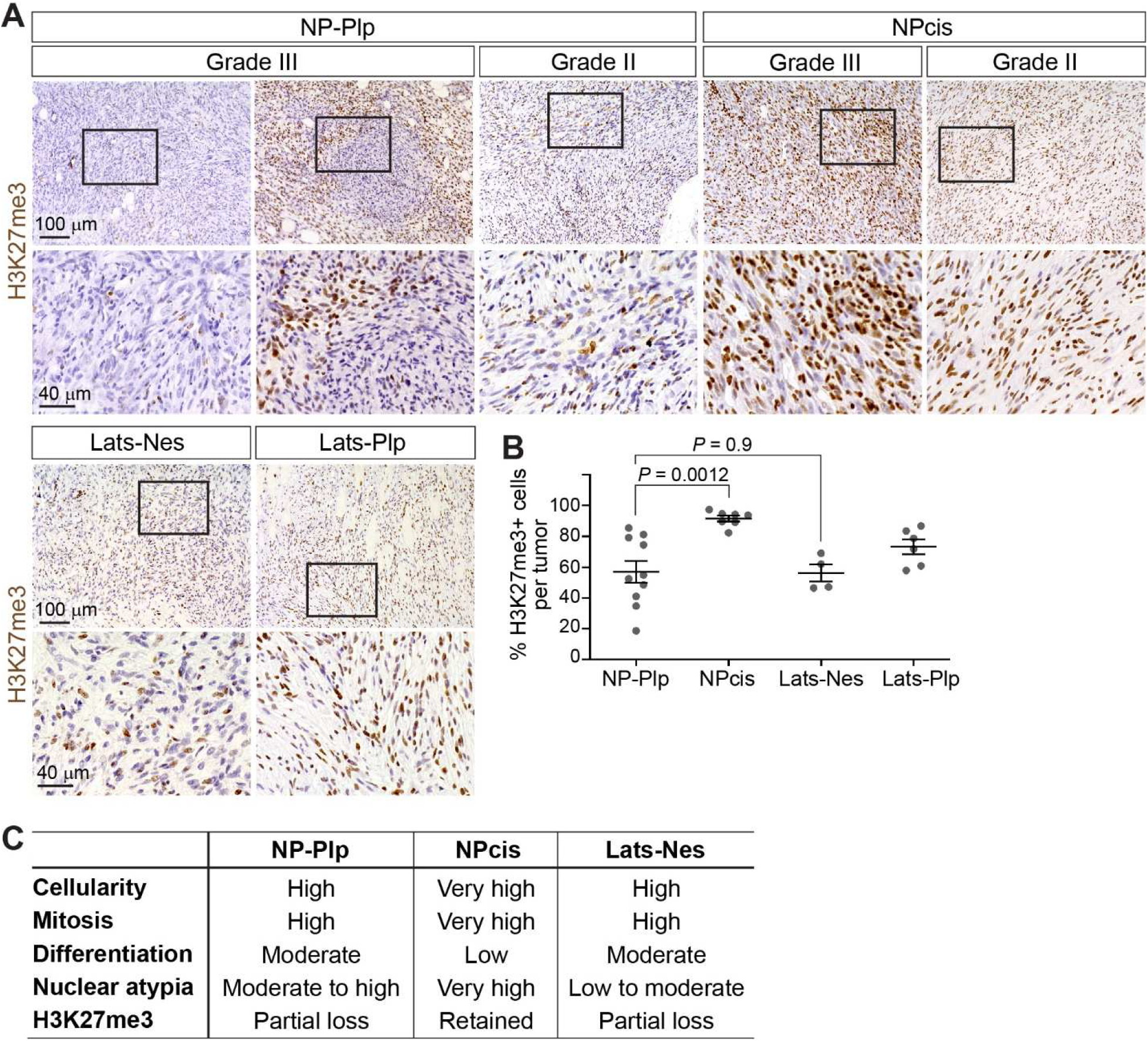
Partial H3K27me3 loss in NP-Plp and Lats GEM-MPNST, but not in NPcis GEM-MPNST. (A, B) H3K27me3 immunostaining and quantification (mean±SEM). Unpaired, two-tailed *t* test. Boxed areas in the upper-row images are enlarged in the lower row. (C) Key histopathologic features of grade III GEM-MPNST in different models.

Figure 4C summarizes the key histopathologic features of grade III GEM-MPNST in these models. NP-Plp and Lats-Nes grade III tumors exhibited very similar features, with the only notable difference being less-frequent nuclear atypia in Lats-Nes tumors. However, NPcis tumors were quite distinct from NP-Plp and Lats-Nes grade III tumors, exhibiting higher cellularity, higher mitotic rates, and more frequent nuclear atypia while being less differentiated.

### *Nf1;p53*-driven and *Lats*-driven GEM-MPNST have distinct transcriptomes

We performed RNA-seq to compare the transcriptomes of NP and Lats tumors. Because many tumors used for RNA-seq analysis lacked matching histologic specimens, we inferred the tumor grade based on observations from histopathologic analyses. For NP-Plp and Lats-Nes tumors, all dermal tumors were inferred as undetermined and all internal tumors as grade III. Only internal NPcis tumors were used for RNA-seq and all were inferred as grade III. All Lats-Plp tumors were inferred as grade II. Both principal component analysis and unsupervised hierarchical clustering segregated NP and Lats tumors (Figure 5A, B). However, these analyses did not segregate tumors of the two NP models or of the two Lats models based on their genotype. Neither did these analyses segregate tumors based on tumor grade, with the caveat that the tumor grades were inferred and were undetermined for many samples. These results suggest that the transcriptomes of NP and Lats tumors are distinct but that the differences between the two NP models and between the two Lats models are small.

**Figure 5.**
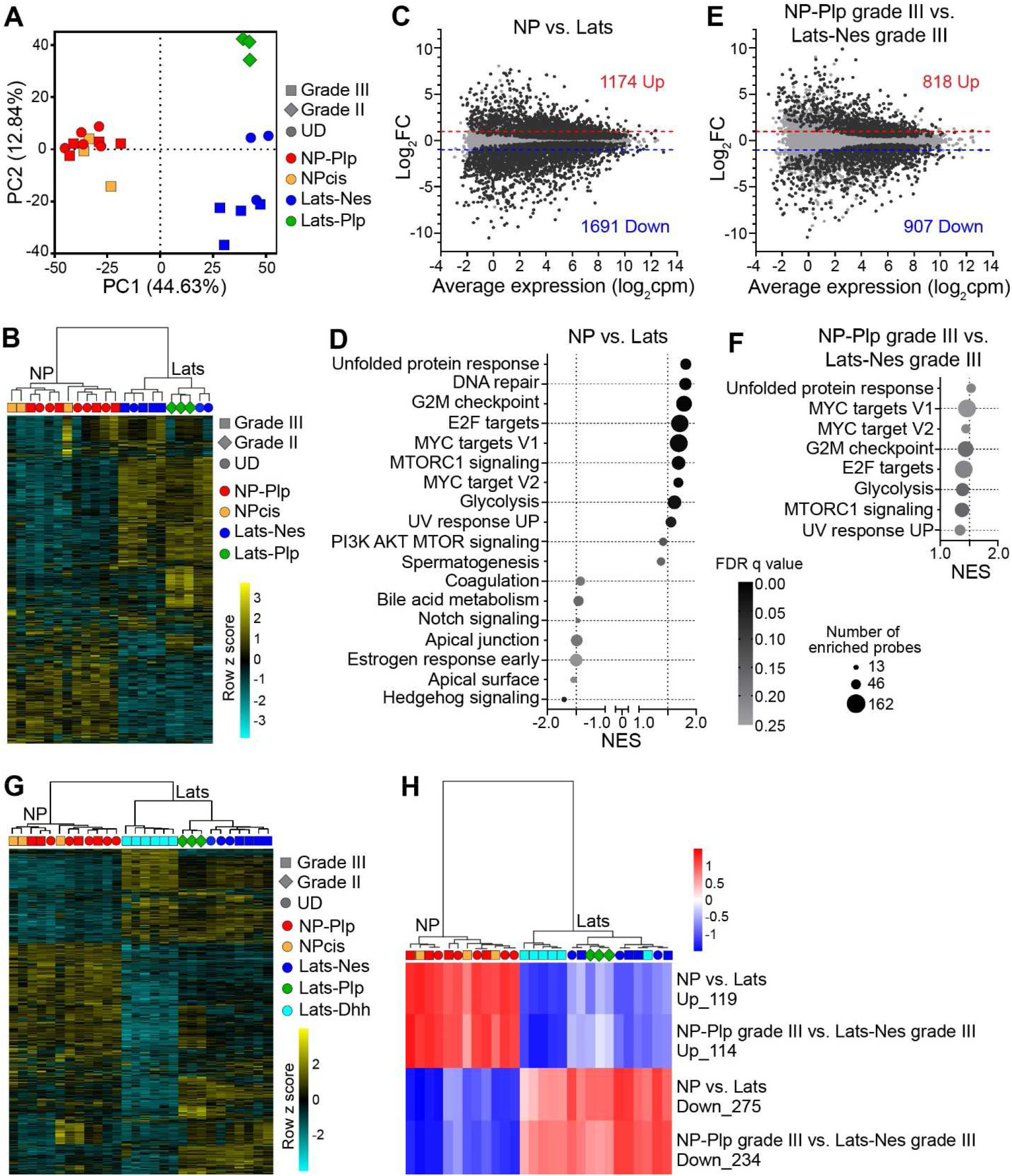
*Nf1;p53*-driven and *Lats*-driven GEM-MPNST have distinct transcriptomes. (A) Principal component analysis of NP and Lats tumors based on the top 3000 most varied genes. UD: tumor grade undetermined. (B) Dendrogram of unsupervised clustering of tumor samples based on the top 2000 most varied genes and the corresponding heatmap. (C) MA plot of differential expression comparing all NP tumors to all Lats tumors. Black dots denote genes with FDR≤0.05. The numbers of significantly up- and down-regulated genes with a fold change (FC) of ≥2.0 or ≤−2.0 are shown. (D) GSEA results showing differentially regulated Hallmark gene sets in NP vs. Lats tumors. (E) MA plot of differential expression comparing grade III NP-Plp to Lats-Nes tumors. (F) GSEA results showing differentially regulated Hallmark gene sets in grade III NP-Plp vs. Lats-Nes tumors. No Hallmark gene sets were upregulated in Lats-Nes tumors. (G) Unsupervised clustering of our NP and Lats tumors and the published Lats-Dhh tumors based on the top 2000 most varied genes. (H) Unsupervised clustering of tumor samples based on single-sample GSEA signatures.

Nearly 3000 genes were significantly (FDR≤0.05) up- or down-regulated with a ≥2-fold change between NP and Lats tumors (Figure 5C). GSEA using the Hallmark gene sets revealed upregulation of gene sets related to growth, proliferation, and cancer, such as the MYC targets, Glycolysis, and PI3K AKT MTOR signaling sets, in NP tumors (Figure 5D). We also compared only grade III NP-Plp tumors to grade III Lats-Nes tumors. Despite being very similar histologically, nearly 2000 genes were significantly up- or down-regulated with a ≥2-fold change (Figure 5E). Most Hallmark gene sets that were upregulated in NP versus Lats tumors were also upregulated in grade III NP-Plp versus grade III Lats-Nes tumors. (Figure 5F). These results further support the conclusion that *Nf1;p53*-driven and *Lats*-driven GEM-MPNST have distinct transcriptomes, and suggest that *Nf1;p53*-driven GEM-MPNST express higher levels of cancer-related genes than do *Lats*-driven GEM-MPNST, even when only grade III tumors are compared.

To confirm that the transcriptomes of NP and Lats GEM-MPNST were distinct, we analyzed the RNA-seq data for grade III Lats-Dhh GEM-MPNST.^12^ Unsupervised clustering clustered Lats-Dhh tumors with our Lats tumors, not with NP tumors (Figure 5G), suggesting that the transcriptome of Lats-Dhh tumors is similar to that of our Lats tumors and distinct from NP tumors. To test this further, we performed ssGSEA, which calculates a separate enrichment score for each pairing of a sample and a gene set, thereby transforming a sample’s gene expression profile to a gene set enrichment profile that represents the activity levels of biological processes and pathways in that sample.^19^ For the gene sets, we used genes that were significantly regulated ≥4-fold in the NP versus Lats comparison or in the grade III tumor comparison with expression levels ≥3 log_2_ counts-per-million. Lats-Dhh tumors again clustered with our Lats tumors, not with NP tumors (Figure 5H). Together, both our own models and published Lats model indicate that *Nf1;p53*-driven and *Lats*-driven GEM-MPNST have distinct transcriptomes.

### *Nf1;p53*-driven GEM-MPNST resemble human MPNST more closely than do *Lats*-driven GEM-MPNST at the transcriptome level

Next, we asked whether *Nf1;p53*-driven or *Lats*-driven GEM-MPNST models were more similar to human MPNST at the transcriptome level. We performed RNA-seq analysis of human MPNST, plexiform neurofibroma (PNF), and neurofibroma (NF) samples. To ensure that our results would be broadly applicable, we included in our analysis published RNA-seq data for human MPNST and PNF samples,^29^ as well as for GEM-neurofibroma from *Nf1*^*fl/fl*^;*Dhh-Cre* mice and paraspinal tumors and GEM-MPNST from *Nf1*^*fl/fl*^;*Ink4a/Arf+/−;Dhh-Cre* mice.^7^ We first transformed the gene expression profile of each sample to ssGSEA signatures, which are less sensitive to technical effects and species differences than are gene-level comparisons. We then took the top 500 most varied gene sets across all mouse tumors as inputs for unsupervised dimensionality reduction of all human and mouse tumors with UMAP (Figure 6A, B). This analysis showed that most *Nf1*-driven GEM-MPNST clustered together, whereas most Lats-driven GEM-MPNST clustered with GEM-neurofibroma from *Nf1*^*fl/fl*^;*Dhh-Cre* mice. Moreover, most *Nf1*-driven GEM-MPNST clustered with the majority of human MPNST samples, whereas Lats-driven GEM-MPNST clustered with most human PNF/NF samples.

**Figure 6.**
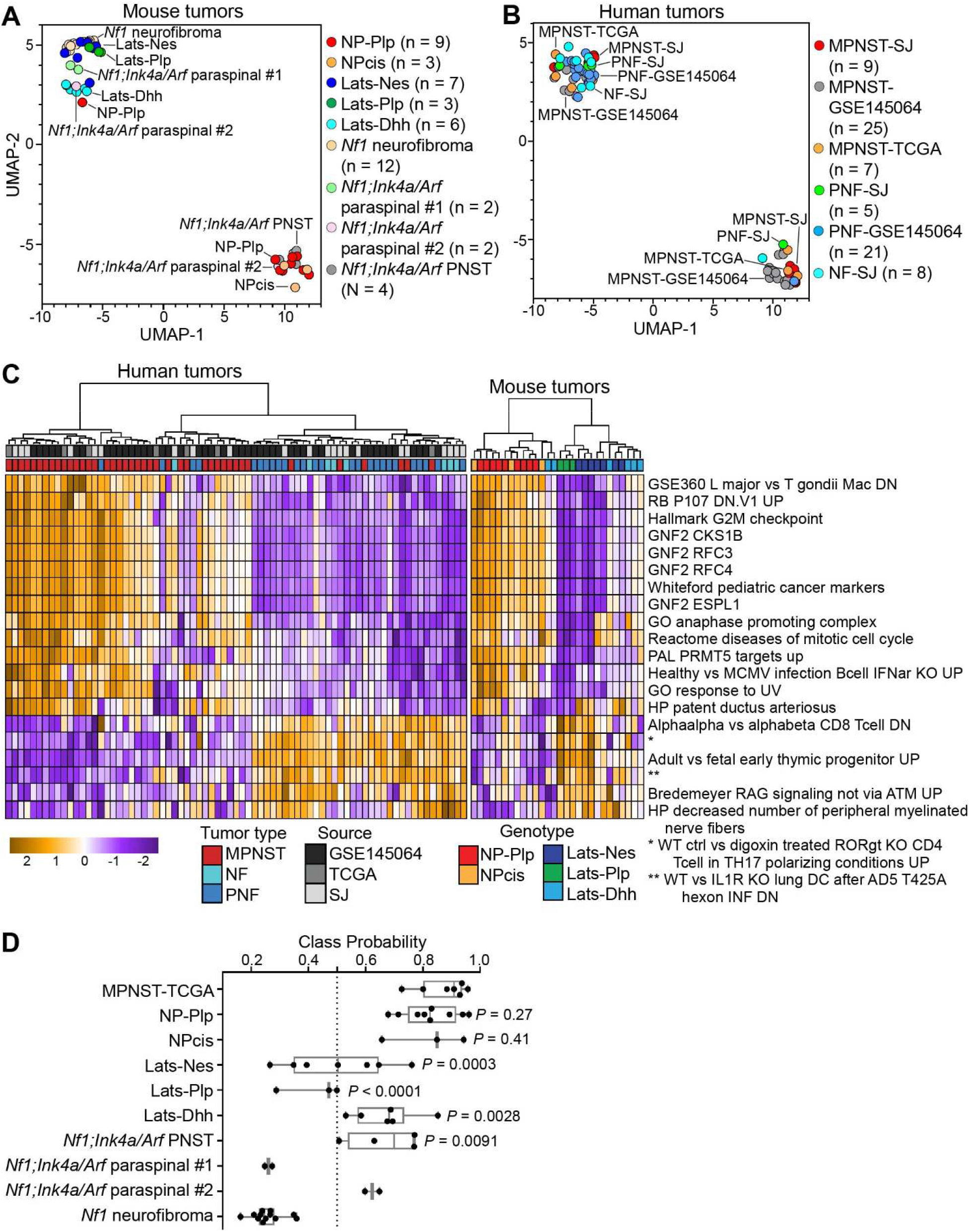
Cross-species transcriptomic comparison of mouse and human nerve sheath tumors. (A, B) UMAP plots of human and mouse tumors based on the top 500 most varied gene sets across all mouse tumors. Mouse and human tumors were analyzed together but are shown on separate plots for visual clarity. MPNST, malignant peripheral nerve sheath tumor. PNF, plexiform neurofibroma. NF, neurofibroma. (C) Unsupervised clustering of tumor samples based on single-sample GSEA signatures of the top 20 gene sets selected by random forest classifier that distinguishes human MPNST from PNF/NF samples. (D) Class probability scores obtained by the random forest classifier. Tumors scoring >0.5 are likely to be MPNST; those scoring <0.5 are likely to be PNF/NF. All comparisons are against MPNST-TCGA. Unpaired, two-tailed *t* test.

To quantify further the similarity between the mouse models and human MPNST, we performed a supervised analysis based on random forest machine learning to discriminate MPNST from PNF/NF samples in our SJ collection and the samples in GSE145064^29^ based on ssGSEA signatures, and used this analysis to derive MPNST class probability scores for the mouse models. The random forest classifier was trained by using the top 20 gene sets that were most discriminative between MPNST and PNF/NF samples (Figure 6C), achieving a test set accuracy of 80%. Using human MPNST samples from TCGA as an independent baseline for comparison, our analysis found that the NP-Plp and NPcis models showed no statistical difference in MPNST class probabilities when compared to TCGA MPNST samples, whereas the other GEM-MPNST models all showed significant differences (Figure 6D). Together, the results of our unsupervised analysis using mouse tumors as the reference and supervised analysis using human tumors as training and testing sets suggest that *Nf1;p53*-driven GEM-MPNST resemble human MPNST at the transcriptome level more closely than do Lats-driven GEM-MPNST.

## Discussion

We established a new mouse model of MPNST by combining an *Nf1;Trp53 cis*-conditional allele and *Plp-CreER*. Postnatal administration of tamoxifen led to GEM-MPNST formation. Our histopathologic and transcriptomic analyses demonstrate that this NP-Plp model closely resembles human MPNST.

The NP-Plp model, by allowing postnatal loss of *Nf1* and *p53*, recapitulates the genetic events in MPNST patients better than existing mouse models. This model also recapitulates the histopathologic features of MPNST better than the NPcis model, which showed only occasional spindle-cell morphology and fascicular and storiform patterns typically found in MPNST.

Inactivation of LATS1/2 of the Hippo pathway was recently shown to cause GEM-MPNST formation.^12^ We made the same discovery independently. Because of the central role of *NF1* in MPNST pathogenesis and the rare occurrence of Hippo pathway mutations in MPNST, we performed detailed comparisons of *Nf1;p53*-driven and *Lats*-driven GEM-MPNST. Although grade III tumors in both the NP-Plp and Lats-Nes models exhibited many histologic features typical of MPNST, nuclear atypia was less frequent in Lats-Nes tumors. Neurofibromin and p53 proteins were lost in NP tumors but preserved in some Lats tumors. However, LATS1/2 protein levels were similarly low in both NP and Lats tumors and TAZ—the transcriptional coactivator suppressed by LATS1/2—was abundant in both models. These results raise the possibility that signaling downstream of LATS1/2 is similarly perturbed in Lats and NP tumors but that signaling downstream of neurofibromin and p53 is perturbed to a greater extent in NP tumors than in Lats tumors. The results of our transcriptomic analysis are consistent with this hypothesis.

Our transcriptomic analysis indicates that *Nf1;p53*-driven and *Lats*-driven GEM-MPNST are molecularly distinct. To determine which GEM-MPNST resembled human MPNST more closely, we performed a cross-species transcriptomic comparison and analyzed our GEM-MPNST, published Lats-Dhh GEM-MPNST^12^ and *Nf1*-driven tumors,^7^ and human tumor samples that we collected or that were published by others.^29^ We took two approaches: one using all of the mouse tumors as a reference to derive the most varied gene sets and clustering the mouse and human tumors based on those gene sets in an unsupervised manner; the other using human samples to train a machine learning classifier that distinguishes MPNST from PNF/NF and scoring mouse tumors with this classifier. Both approaches indicated that *Nf1;p53*-driven GEM-MPNST resemble human MPNST more closely than do *Lats*-driven GEM-MPNST.

Complete and partial H3K27me3 loss is common in MPNST.^30^ NP-Plp and Lats GEM-MPNST, but not NPcis GEM-MPNST, exhibited partial H3K27me3 loss. The mechanism underlying H3K27me3 loss is unclear; we detected no significant reduction in *Suz12*, *Eed*, or *Ezh1;2* mRNA levels in NP-Plp tumors as compared to NPcis tumors, nor consequential changes in their mRNA sequences. Recent studies found H3K27me3 loss in many types of solid tumors, most of which might not be caused by PRC2 inactivation.^30^

In summary, we have shown that the NP-Plp GEM-MPNST model resembles human MPNST genetically, histologically, and molecularly—more so than the NPcis model and *Lats*-driven GEM-MPNST models. The NP-Plp model is genetically simple, requiring only two modified alleles, making it easy to maintain and an ideal platform for preclinical studies. Given its tamoxifen-inducible nature, this model can be used to study the time/stage dependency of the tumorigenic potential of Schwann cells.

## Supporting information

List of primary antibodies

## Acknowledgements

We thank the Veterinary Pathology core, the Hartwell Center, the Cell and Tissue Imaging Center, Cytogenetics, and Biorepository at St. Jude Children’s Research Hospital for assistance with data acquisition and analysis, Dr. Alfonso Lavado for assistance with mouse maintenance, and Dr. Keith A. Laycock for scientific editing of the manuscript.

